# Diminishing returns of inoculum size on the rate of a plant RNA virus evolution

**DOI:** 10.1101/226357

**Authors:** Rebeca Navarro, Silvia Ambrós, Fernando Martínez, Santiago F. Elena

## Abstract

Understanding how genetic drift, mutation and selection interplay in determining the evolutionary fate of populations is one of the central questions of Evolutionary Biology. Theory predicts that by increasing the number of coexisting beneficial alleles in a population beyond some point does not necessarily translates into an acceleration in the rate of evolution. This diminishing-returns effect of beneficial genetic variability in microbial asexual populations is known as clonal interference. Clonal interference has been shown to operate in experimental populations of animal RNA viruses replicating in cell cultures. Here we carried out experiments to test whether a similar diminishing-returns of population size on the rate of adaptation exists for a plant RNA virus infecting real multicellular hosts. We have performed evolution experiments with tobacco etch potyvirus in two hosts, the natural and a novel one, at different inoculation sizes and estimated the rates of evolution for two phenotypic fitness-related traits. Firstly, we found that evolution proceeds faster in the novel than in the original host. Secondly, we found the predicted diminishing-returns effect of inoculum size on the rate of evolution for one of the fitness traits, but not for the other, which suggests that selection operates differently on each trait.

## Introduction

An essential question in Evolutionary Biology is how mutation rate, population size, the magnitude of beneficial mutational effects, and the load of deleterious mutations interact to determine the rate at which asexual populations evolve. Laboratory experiments conducted with viruses [1,2], bacteria [3–5] and yeast [6] have shown that, above certain limits, population size has a diminishing-returns effect on the rates of evolution, meaning that there is little gain in the rate of evolution by increasing populations size and mutation rates (*i.e*., the supply of beneficial mutations). This phenomenon is known as clonal interference [7–9]. Clonal interference has been shown also to play a significant role during the epidemiological spread and antigenic diversification of *Influenza A virus* [10]. Essentially, clonal interference means that beneficial alleles in coexisting lineages within a large population must compete each other in their way to fixation and, thus, only the best one of them will prevail. Afterwards, the second allele will appear and rise its frequency to fixation but in the genetic background of the first one (assuming it is still beneficial in it). Therefore, in the clonal interference regime beneficial alleles will fix sequentially. The rate at which beneficial alleles fix in the population is given by

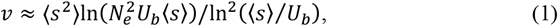

where 〈*s*〉 is the average beneficial fitness effect, *U_b_* the rate at which beneficial mutations are produced and *N_e_* the effective population size [8]. Indeed, one of the advantages of sexual reproduction is pooling together into the same genome both beneficial alleles, thus relaxing clonal interference. In small populations, beneficial alleles will also fix sequentially but for a different reason: because the probability of appearing beneficial mutations and survive drift while they are infrequent is low [7–9]. The rate of evolution in such a successional-mutation regime is then given by the simpler relationship

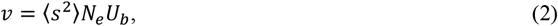

which indicates that the rate of evolution should increase linearly with the availability of beneficial alleles *N_e_U_b_* and their average square beneficial fitness effect.

Population bottlenecks are pervasive events during virus transmission from host to host but also between different tissues within an individual infected host [11]. Very severe bottlenecks turn on Muller’s ratchet, a process that results in an increase in the genomic load of deleterious mutations, with a concomitant decline in fitness [12–14] that theoretically may drive populations to extinction [15]. Indeed, it has been shown for *Vesicular stomatitis virus* that the size of the bottleneck leading to the onset of Muller’s ratchet depends on the fitness of the genotype used in the experiments and on the standing beneficial genetic variation present in the inoculum [16–18]. In large enough populations, such variation quickly amplifies and wipes out deleterious variants, thus slowing down and even reverting the fitness decay process.

Though in general one viral infectious unit is enough to trigger infection [19,20], many relevant properties of infection, such as the total amount of virus accumulated, the immune response from the host and the severity of symptoms directly depend on the inoculum size [21,22].

So far, most of the studies mentioned exploring the interplay between *N_e_*, *U_b_* and *v* have been performed in oversimplified experimental systems such as cell cultures [1,2,12–18] or *in vitro* with artificial media [3–6]. With this study, we want to expand this observations to a biologically fully realistic experimental system: the pathosystems formed by *Tobacco etch virus* (TEV; genus *Potyvirus*, family *Potyviridae)* and its natural host tobacco (*Nicotiana tabacum* L) and novel host pepper *(Capsicum annuum* L). After assessing the number of TEV RNA genomes per infectious unit in both hosts, we performed evolution experiments by serial mechanical passages at different inoculation doses (N). Then we evaluated two TEV fitness-related traits along the experimental passages (viral load and infectivity) and estimated their rates of evolution.

## Experimental methods

### Virus, plants and growth conditions

Plasmid pMTEV contains the TEV genome from a tobacco isolate [23]. A stock of infected tissue was generated before starting the evolution experiment as described elsewhere [24]. Host species *N. tabacum* cv. Xanthi and *C. annuum* cv. Marconi (both from the *Solanaceae* family) were used as experimental hosts. In both, TEV produces systemic symptoms of different severity. For all the experimental steps, plants were maintained in a BSL2 greenhouse at 25 °C and a 16 h photoperiod.

### Estimating infectivity

The infectivity of each viral sample along the evolution experiments was evaluated as the frequency of plants showing symptoms of infection between 9 and 14 days post-inoculation (dpi). The LaPlace’s estimator for the Binomial parameter 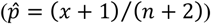 was used instead of the classic maximum-likelihood estimator because it provides more robust and reliable estimates for small sample sizes, *n* [25].

### Estimating viral load

Ground material was obtained from the whole infected plant after removing the inoculated leaf. Total RNA extraction from 100 mg of grounded tissue per plant was performed using Plant Isolation RNA Mini Kit (Agilent) following the manufacturer’s instructions. The concentration of total RNA in the extracts was spectrophotometrically determined using a NanoDrop ND100 (Thermo Scientific) and adjusted to 40 - 55 ng/μg for each sample. Each of these normalized RNA extracts was used to quantify TEV concentration by RT-qPCR using primers and methods previously described [26]. Amplifications were done with a OneStep Plus Real-Time PCR System (Applied Biosystems) according to the following profile: 5 min at 42 °C, 10 s at 95 °C following 40 cycles of 5 s at 95 °C and 34 s at 60 °C and melt curve stage or 95 °C 15 s, 60 °C 1 min and 95 °C 15 s. RT-qPCR reactions were performed in triplicate for each sample. Quantification results were further examined using the StepOne Software v.2.2.2 (Applied Biosystems). The resulting measures have units of TEV genomes per ng of total RNA.

### Evaluating the number of genomes per infectious unit

The data for tobacco and pepper used for these analyses were taken from [19]. In short, in this previous study genetically engineered variants of TEV expressing either of two fluorescent proteins (eGFP and mCherry) were used to identify and count infection foci (by definition, each produced by a single infectious unit). The viruses were serially diluted and each dilution inoculated into either host plants (4 replicates each). Plants were observed daily with a Leica MZ16F stereomicroscope, using a 0.5 × objective lens, and GFP2 and DSR filters (Leica) to view eGFP and mCherry respectively. In parallel, a fraction of each one of the dilutions was used to quantify the number of viral genomes by RT-qPCR as described above.

An additional similar experiment was performed with the wildtype TEV but inoculating quinoa (*Chenopodium quinoa* Willd) leaves. In this host, TEV generates local necrotic lesions that are readily visible.

### Experimental evolution

Three evolutionary regimes were defined that differed in the inoculum size; that is, in the availability of potential beneficial mutations present in the inoculum, *N* ∝ *N_e_U_b_*. The first treatment corresponded to the inoculum resulting from adding 500 μl of inoculation buffer (50 mM potassium phosphate pH = 7.0, 3% PEG, 10% Carborundum) to 500 mg of infected tissue homogenized with liquid N_2_ in a mortar with a pestle; hereafter referred as the 10N-inoculum size treatment. The second treatment consisted in adding 900 μl of inoculation buffer to 100 mg of homogenized tissue, and will be referred as the *N*-inoculum size treatment. For the third treatment, we made a 1:10 dilution from the *N*-inoculum size by adding 900 μl of inoculation buffer to 100 μl of this extract, and we will refer to this treatment as the *N*/10-inoculum size. Ten μl were inoculated on each plant except in the case of the 10N-inoculum size, in which two leafs were inoculated with 1C μl each.

Five independent evolution lineages for each of these three treatments and host species were founded and maintained by 15 serial passages. Fourteen dpi, the aerial part of each plant was collected, homogenized, diluted as needed and used to inoculate the next batch of plants. This protocol was repeated a total of 15 times (see [24,27-29] for a more detailed description of serial passages evolution experiments).

### Statistical analyses and inferences of the rates of evolution of fitness-related traits

Viral load and infectivity, two essential fitness components, were measured at passages 0, 3, 5, 8, 11, 13, and 15. The resulting time-series data were then fitted to first-order autoregressive integrated moving average models, ARIMA(1,0,0), of the form

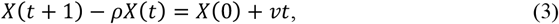

where *X* represents the value of the fitness trait being analyzed at time *t*, *ρ* measures the correlation between observed values at two consecutive time points (self-similarity in the time series) and *v* corresponds to the net rate of change of the fitness trait, *i.e.*, the rate of evolution. Since we are interested in exploring the effect of *N* into *v*, we have estimated an independent *v* value for each of the five lineages evolved under each of the three evolutionary regimes. These data were then fitted to the following factorial ANOVA model:

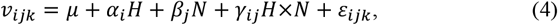

in which *H* stands for the two host species used in the evolution experiment (tobacco and pepper) and *N* for the three inoculum sizes tested. These two factors were considered as orthogonal. Finally, *μ* represents the grand mean value and *ε_ijk_* the error associated to each individual measure and assumed to be Gaussian.

Other statistical tests used will be introduced as needed along the next section.

## Results

### The number of genomes per infectious unit varies among host species

Fig. 1 shows the observed relationship between the number of TEV genomes (gRNA) and the number of infectious units (*LFU*). The relationship is linear in the log-log scale. Data were fitted to the following linear model by means of GLMM technique:

**Fig. 1:**
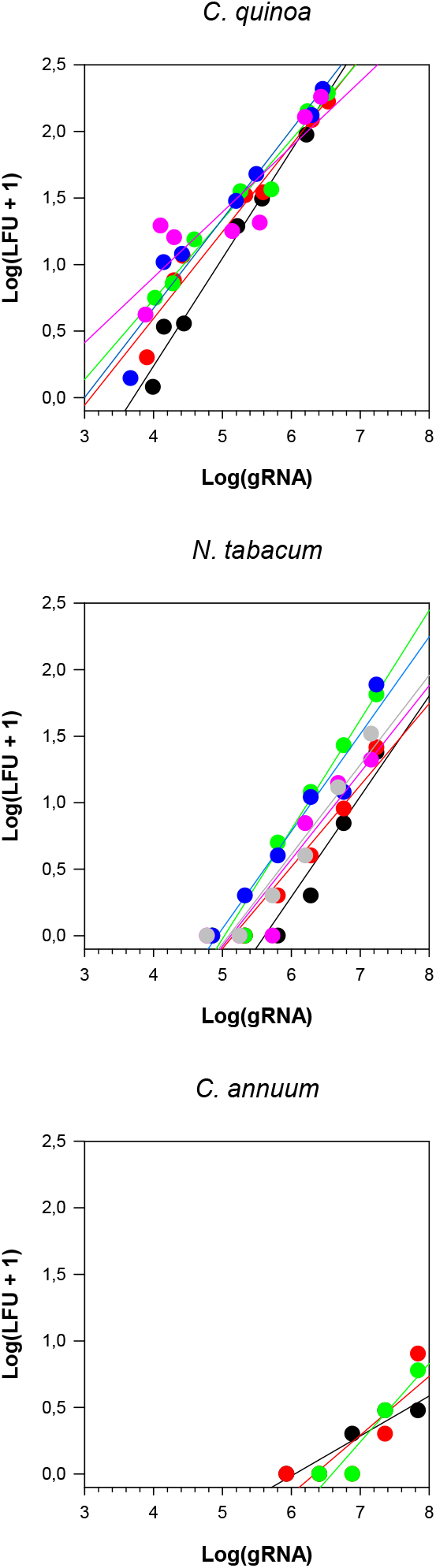
Relationship between TEV genomes (gRNA) and infectious units (*LFU*). Different colors represent different replicates of the experiment. Each panel corresponds to the indicated host species.

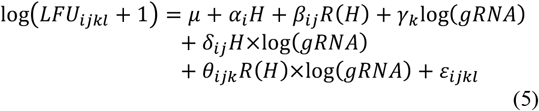

where, as above, factor *H* stands for the three host species used in these experiments (quinoa, tobacco and pepper), factor *R* for the biological replicates of the experiment done for each host (*i.e*., *R* is nested within *H*), and log(*gRNA*) is incorporated into the model as a covariable. The model also includes the interactions between the covariable and the two factors. *μ* and *ε_ijkl_* have the same meaning than in eq. (4). The significance of each factor in the model was assessed by means of likelihood ratio tests.

Net differences among host species exist (*χ*^2^ = 36.640, 2 d.f., *P* < 0.001), indicating that, on average, different numbers of TEV gRNAs are needed to initiate an infection, with quinoa being the host that requires the less and pepper requiring the most. Obviously, the covariable has a net effect in the number of infectious units (*χ*^2^ = 163.013, 1 d.f., *P* < 0.001): the more TEV gRNAs are inoculated, the more infection foci are produced. *R* had no significant effect on the number of infectious units by itself nor in the interaction with the covariable (in both cases, *P* ≥ 0.084), indicating the high reproducibility of the results. More interestingly, the slope of the regression lines varies among host species (significant interaction term between *H* and the covariable: *χ*^2^ = 10.650, 2 d.f., *P* = 0.005), indicating that equivalent increases in the gRNA dosage do not result in similar increases in the number of infectious units on each host. On average, slopes are flater for pepper, indicating that more gRNAs have to be inoculated in order to achieve the same number of infectious units, and steaper for tobacco and quinoa (no differences among them: Tukey *post-host* test, *P* > 0.05). Indeed, 30.4 more TEV gRNAs have to be inoculated into pepper in order to generate an equivalent number of infectious units than in tobacco.

Therefore, from these experiments, we concluded that for the evolution experiments to be performed in equivalent conditions of *N* in both experimental hosts, it is necessary to add ~30 times more TEV gRNA to pepper than to tobacco.

### Inoculum size has a diminishing-returns effect on the rate of evolution of fitness traits

Supplementary fig. S1 contains the raw time series data for the two fitness components, viral load and infectivity. The data for each evolutionary lineage were fitted to the model shown in eq. (3) to generate independent estimates of v. Fig. 2 shows the average estimates of these rates of evolution for each fitness trait, experimental host and inoculum size.

**Fig. 2:**
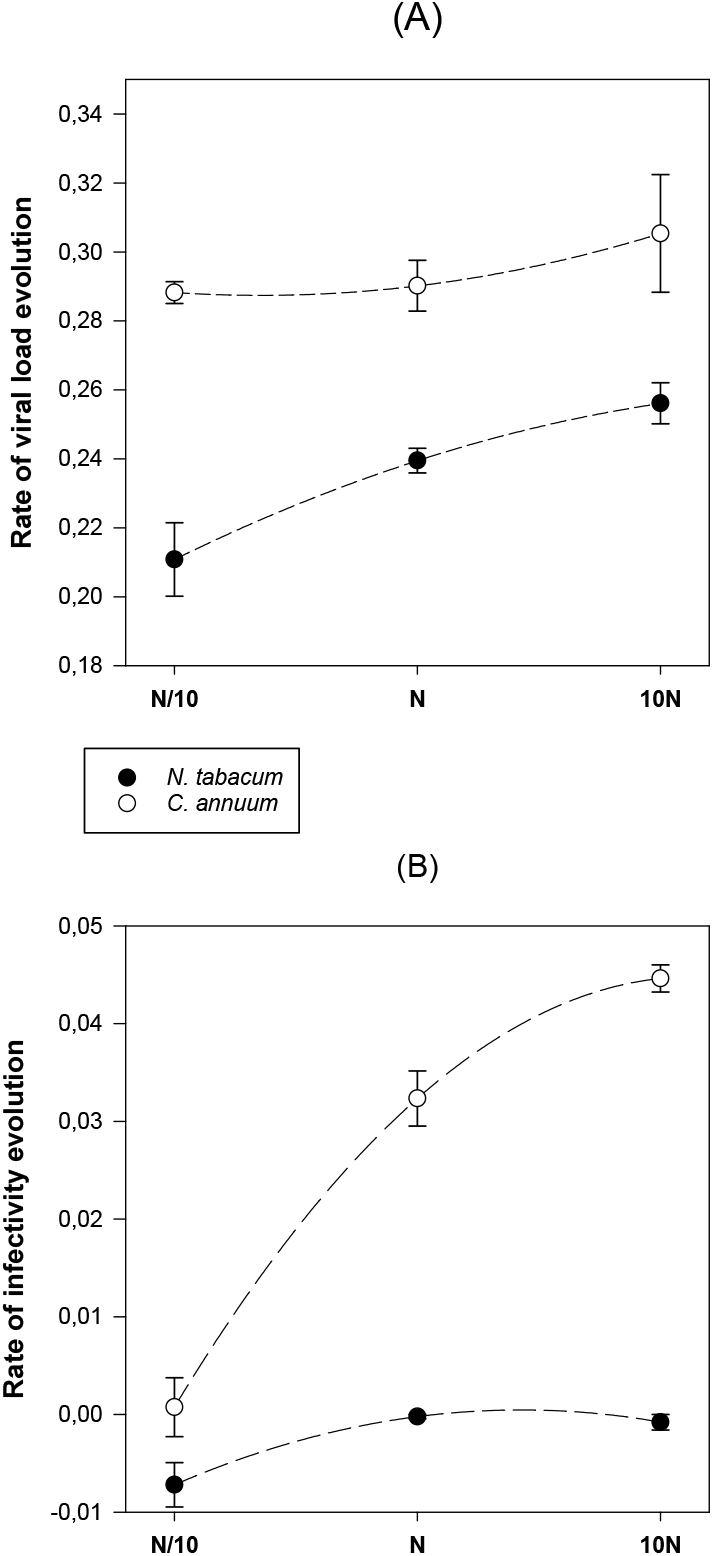
Effect of host species and inoculum size in the rate of evolution of two fitness traits, viral load (A) and infectivity (B). Error bars represent ±1 SD.

Focusing first into the viral load, data shown in fig. 2A were fitted to the ANOVA model described by eq. (4). A highly significant effect was associated to the *H* factor (*F*_1,24_ = 60.812, *P* < 0.001). On average, rates of evolution were higher in the novel host pepper than in the natural reservoir tobacco. More interestingly, *N* also had an overall significant effect on the rates of evolution (*F*_2,24_ = 5.652, *P* = 0.010), with slower rates being associated to smaller inoculum sizes (*N*/10) and increasing in magnitude as *N* increases. No significant interaction between *H* and *N* has been found (*F*_2_,_24_= 1.457, *P* = 0.253), thus suggesting the effect of *N* on *v* was similar for both experimental hosts.

Moving our attention now on the second fitness trait, infectivity, the data shown in fig. 2B were also fitted to the ANOVA model shown in eq. (4). *H* had a highly significant effect (*F*_1,24_ = 297.719, *P* < 0.001), again with lineages evolved in the novel host characterized by faster evolutionary rates than lineages evolved in the natural host. *N* also had a significant overall positive effect on *v* (*F*_2,24_ = 87.751, *P* < 0.001), with rates increasing in magnitude with *N*. In the case of infectivity, a significant *H*×*N* has been found (*F*_2,24_ = 43.884, *P* < 0.001), with increases in *N* having larger effects in the novel host than in the natural one.

The clonal interference hypothesis discussed in the Introduction predicts a diminishing-returns effect of the amount of beneficial diversity contained in the inoculum (*N* in our notation) on the rate of adaptation *v*. To test this hypothesis, we fitted a linear and a quadratic model to each dataset in fig. 2, as done in [1]. The linear model representing the null hypothesis of *v* being directly proportional to the availability of beneficial variation as described by eq. (2) and the quadratic model illustrating the diminishing-returns effect (a downwards curvature with increasing *N*). In the case of viral load (fig. 2A), the null hypothesis could not be rejected for either host according to a partial-*F* test comparing the goodness-of-fit of both nested models (*F*_1,12_ = 0.450, *P* = 0.515 for tobacco and *F*_1,12_ = 0.245, *P* = 0.630 for pepper). By contrast, in the case of infectivity (fig. 2B), the null hypothesis was rejected in both cases, as the goodness-of-fit significantly improved by adding a quadratic term with negative coefficient (*F*_1,12_ = 4.800, *P* = 0.049 for tobacco and *F*_1,12_ = 9.719, *P* = 0.009 for pepper).

Therefore, from these experiments we conclude that rate of adaptation depends on both the host species (being faster in novel hosts) and the inoculum size. However, the clonal interference effect could only be detected for one of the fitness traits, infectivity, but not for viral load.

## Discusion

The results here reported confirm the role of clonal interference in the evolution of asexual RNA viruses and expand the observation from a simple animal cell culture systems [1,2] to a complex biologically realistic plant pathosystem, in which multiples sources of selection may be operating simultaneously over the virus population. As expected from their large population size and high genomic mutation rates (in the case of TEV in the range 0.01 – 0.1 per genome and replication event [30]), the number of possible beneficial mutations coexisting at a given time in a population must be certainly large, as shown recently by ultra-deep sequencing of evolving TEV lineages [24].

One of our more relevant observation is that the rate of adaptation was significantly faster in the novel host than in the natural reservoir. This is expected as more beneficial variation may exist in a novel host, whereas in the reservoir host this variation may have been already exhausted. Supporting this notion, Cuevas *et al*. [24] found that the temporal evolution of TEV genetic variability within the reservoir host was consistent with the presence of a dominant genotype and the steady accumulation of neutral alleles. By contrast, in pepper, at least in one of the experimental replicates, the ancestral virus was quickly eliminated by a clone carrying a beneficial mutation. In its way to fixation, this clone also displaced another beneficial mutation that was later on fixed on the new dominant genetic background [24].

Why clonal interference seems to be more important for infectivity than for viral load? A plausible explanation is that more beneficial genetic variation exists to improve infectivity, perhaps because it is a genetically simpler trait: it only requires entering into the cell and being able of initiating infection. By contrast, viral load, which relates to within cell replication, cell-to-cell and systemic movement and virion stability, is a more complex trait which requires from the intervention of multiple host factors at different stages. This complexity makes reasonable to think that, given the compactness of viral genomes and multifunctional activities of their proteins, beneficial mutations improving one particular interaction perhaps might have a negative pleiotropic effect on other interactions, thus making the number of possible beneficial mutations smaller and therefore clonal interference weaker for viral load.

It has been theorized that transmission bottlenecks may provide advantages to RNA virus populations. One of such advantages being to facilitate moving from adaptive peaks in a rugged fitness landscapes. On such landscapes, it becomes readily possible for the virus to become trapped on suboptimal fitness peaks [31], and transmission bottlenecks may allow genetic variants to get fixed into distant regions of the landscape. Another possible advantage of transmission bottlenecks may be to remove cheaters (e.g., defective interfering viruses) from the population [32]. Widespread transmission bottlenecks are likely one important reason why interfering viruses are not so common in natural pathosystems.

We thank F. de la Iglesia and P. Agudo for excellent technical support. This work was supported by grant BFU2015-65037-P from Spain’s Ministry of Economy, Industry and Competitiveness and by the Santa Fe Institute.

